# Genetic background influences the 5XFAD Alzheimer’s disease mouse model brain proteome

**DOI:** 10.1101/2023.06.12.544646

**Authors:** Cheyenne D. Hurst, Amy R. Dunn, Eric B. Dammer, Duc M. Duong, Nicholas T. Seyfried, Catherine C. Kaczorowski, Erik C. B. Johnson

## Abstract

There is a pressing need to improve the translational validity of Alzheimer’s disease (AD) mouse models. Introducing genetic background diversity in AD mouse models has been proposed as a way to increase validity and enable discovery of previously uncharacterized genetic contributions to AD susceptibility or resilience. However, the extent to which genetic background influences the mouse brain proteome and its perturbation in AD mouse models is unknown. Here we crossed the 5XFAD AD mouse model on a C57BL/6J (B6) inbred background with the DBA/2J (D2) inbred background and analyzed the effects of genetic background variation on the brain proteome in F1 progeny. Both genetic background and 5XFAD transgene insertion strongly affected protein variance in hippocampus and cortex (n=3,368 proteins). Protein co-expression network analysis identified 16 modules of highly co-expressed proteins common across hippocampus and cortex in 5XFAD and non-transgenic mice. Among the modules strongly influenced by genetic background were those related to small molecule metabolism and ion transport. Modules strongly influenced by the 5XFAD transgene were related to lysosome/stress response and neuronal synapse/signaling. The modules with the strongest relationship to human disease—neuronal synapse/signaling and lysosome/stress response—were not significantly influenced by genetic background. However, other modules in 5XFAD that were related to human disease, such as GABA synaptic signaling and mitochondrial membrane modules, were influenced by genetic background. Most disease-related modules were more strongly correlated to AD genotype in hippocampus compared to cortex. Our findings suggest that genetic diversity introduced by crossing B6 and D2 inbred backgrounds influences proteomic changes related to disease in the 5XFAD model, and that proteomic analysis of other genetic backgrounds in transgenic and knock-in AD mouse models is warranted to capture the full range of molecular heterogeneity in genetically diverse models of AD.

## Introduction

Alzheimer’s disease (AD) is the leading cause of dementia worldwide, with limited treatment options currently available. Clinical trials of potential disease-altering therapies for AD have had very high failure rates. Evaluation of failings at the preclinical stage have implicated poor translation between widely-used mouse models of AD and the human disease as a potential explanation (1). Murine models of AD represent a cornerstone of disease research and have played a critical role in the preclinical development process. Traditional modeling of AD in mice has primarily focused on the overexpression of the amyloid precursor protein (APP) and presenilins 1 and 2 (PSEN1/2) that harbor familial AD (FAD) mutations. Such transgenic FAD models robustly develop β-amyloidosis and have been instrumental in understanding disease mechanisms and progression (2). However, no single model fully recapitulates the complex pathobiology of human disease, leading to gaps in cross-species translation. The need to develop and improve the validity and translatability of AD mouse models remains an ongoing challenge.

Mouse models of AD have been almost exclusively developed on a single inbred strain, the C57BL/6J (B6). The selection of B6 as the ideal background for developing AD model systems has been largely due to their desirable behavioral characteristics and the development of cognitive impairment and amyloid deposition (3–6). For instance, the commonly used 5XFAD transgenic model was developed on the B6 background to overcome variable phenotypes produced on a hybrid background (7). However, the utility of inbred strains has come under question in recent years as research has indicated the lack of generalizability of, and often opposing, phenotypes across different strains expressing FAD mutations. Outcome measures of interest to AD researchers, such as learning and memory specific task performance and amyloid pathology, are significantly impacted by mouse genetic background (8–10).

Genetics contributes strongly to susceptibility and person-to-person heterogeneity for both familial and sporadic forms of AD. Twin studies have estimated the heritability of AD to be approximately 60-80% (11). Even among autosomal dominant cases, onset of clinical symptoms can vary across families that carry the same mutation (12). Together, these findings highlight the complex interaction between genetic factors and phenotype in human AD and indicate that there are undefined genetic factors that contribute to disease risk or resilience. The complex relationship between genetic composition and phenotypic variability observed in human populations, of which is lacking within genetically homogenous inbred mouse strains, has led to the hypothesis that incorporating genetic variability in AD mouse models could improve translatability. In support of this hypothesis, an AD transgenic mouse reference panel (AD-BXD) was recently developed to explore the impact of genetic complexity on phenotypic segregation and translation with human disease features (13). The reference panel was generated by crossing 5XFAD mice on a B6 background with the BXD recombinant inbred strain series derived from B6 and DBA/2J (D2) crosses. Results from this study identified improved AD phenotypes relating to varied age of onset and rate of memory impairment across the resulting strains. These findings indicate high potential value for incorporating genetic variability as a means to improve the face validity of transgenic mice. However, it is currently unknown how such genetic diversity influences protein changes related to disease in such models.

Quantitative proteomics has contributed to improved understanding of human brain pathophysiology in AD (14, 15). Here we asked how genetic diversity introduced by crossing B6 and D2 mice (B6xD2), two fully inbred strains, affects the mouse brain proteome and its perturbation by the 5XFAD transgene. We found strong proteomic alterations related to both genetic background and transgene expression. We then integrated human brain proteomic data to assess relevant overlap and cross-species relationships. Our findings serve as a resource for the continued development of AD mouse models with improved translational power.

## Results

### Genetic background and AD genotype strongly contribute to proteomic variance

To address how genetic variation in AD mouse model systems impacts the brain proteome, label free quantitation mass spectrometry (LFQ-MS) analysis was performed on brain tissue from two brain regions (cortex and hippocampus) from B6 and B6xD2 mouse lines as previously described (16). The cortex and hippocampus are both vulnerable in AD and represent relevant regions of interest for proteomic analysis. Animals were aged 13-14 months, a time at which significant pathology has developed in the 5XFAD model, and balanced by sex (7, 17). Protein abundance data were pre-processed to remove proteins with higher missing values and sample outliers, resulting in a final data matrix of 3,368 proteins measured across 70 total samples (**Fig. 1; Supplemental Tables 1-2**). The term “line” will be used throughout to refer to the mouse genetic background (i.e. B6 or B6xD2) while “AD genotype” is used to denote the transgenic manipulation of the recipient mouse (Ntg vs 5XFAD).

**Figure 1:**
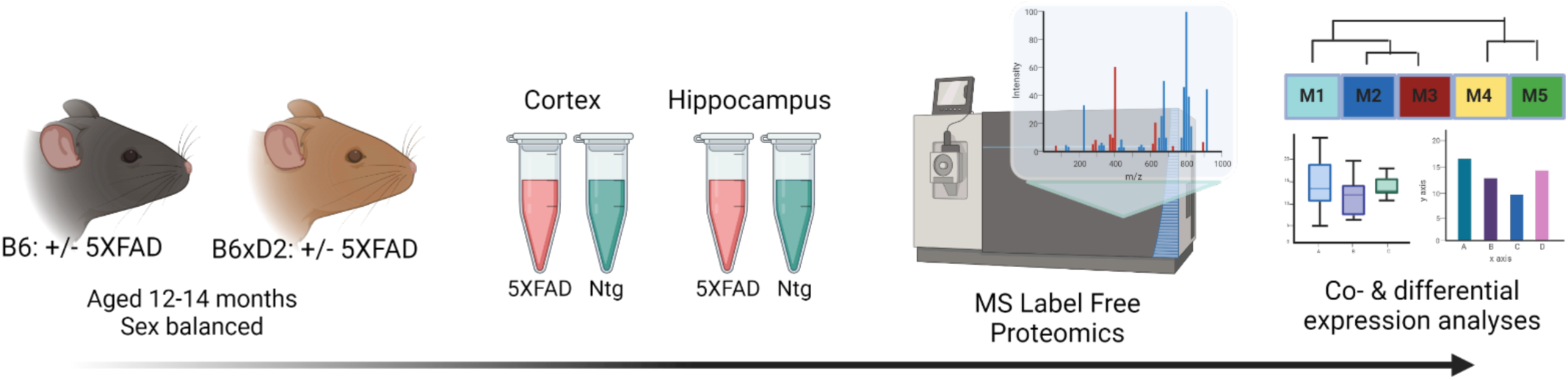
Study overview. Schematic representation of experimental workflow for matched mouse brain tissue samples from cortex and hippocampus from 5XFAD and nontransgenic (Ntg) mice on B6 (n=10 5XFAD, n=10 Ntg) or B6xD2 (n=10 5XFAD, n=10 Ntg) genetic backgrounds (n=40 total per region). Tissue samples underwent enzymatic digestion and label free mass spectrometry analysis followed by co- and differential expression analyses.

To assess the primary contributing features of variance within the proteome, we calculated the top five principal components (PCs) of the data for each brain region and correlated the PCs with line to assess genetic background effects, AD genotype to assess transgene effects, and sex (**Fig. 2A, B**). AD genotype strongly correlated with the first PC in both brain regions. More variance was explained by AD genotype in hippocampus (23%) than cortex (16%). Line significantly correlated with more than one PC in each brain region, indicating that genetic background had broad effects on protein variance across the proteome. Variance partition analysis verified line and AD genotype as the largest drivers of variation in protein abundance in each brain region (**Fig. 2C, D; Supplemental Tables 3-4**). To identify proteins with the highest total variance attributable to each trait, variance partition results were sorted and ranked by protein for both cortex and hippocampus (**Fig. S1A, B**). Many proteins in the top 10 variance for each trait overlapped between brain region. Among the proteins with the highest attributable variance to line, proteins with significantly higher or lower abundance in B6xD2 compared to B6 could be observed, as expected (**Fig. S1C**). Conversely, those proteins with the highest attributable variance to AD genotype were not significantly different between genetic backgrounds (**Fig. S1D**). These results suggested that genetic background variation had a large effect on the mouse brain proteome in both hippocampus and cortex. In cortex, the effect was comparable to that observed with 5XFAD transgene expression, whereas in hippocampus, the effect of AD genotype on protein expression variance was greater than the effect of genetic background.

**Figure 2:**
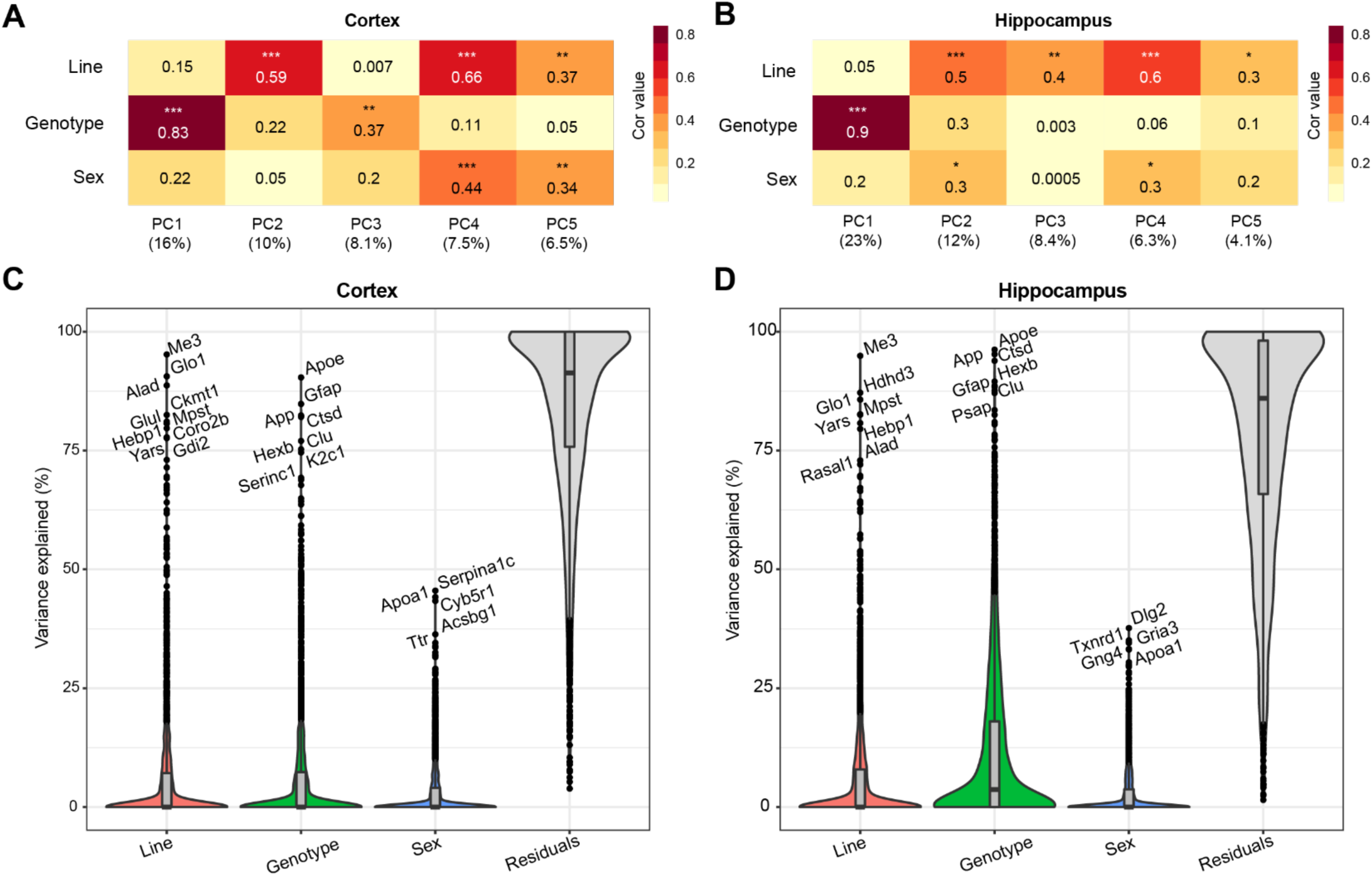
Genetic background and AD genotype strongly contribute to proteomic variance. **A-B**) Principal component analysis (PCA) in cortex (**A**) and hippocampus (**B**) was used to examine the relationship between the top five PCs and line, AD genotype and sex traits using Spearman’s rho correlation. Heatmaps represent the strength of correlation between traits and PCs (darker color indicates increasing strength of correlation). AD genotype was the most strongly associated trait with PC1 in the cortex (ρ= 0.83) and hippocampus (ρ= 0.9). *P* *<0.05, **<0.01, ***<0.001. **C-D**) Variance partition analysis was performed for cortex (C) and hippocampus (**D**) to quantify overall variance explained by each trait. Line and AD genotype contributed to the highest percent variance explainable in both brain regions.

### Protein abundances driven by AD genotype and genetic background

To examine how protein abundances were impacted by either AD genotype or genetic background, we assessed differential protein abundance in each brain region. For AD genotype, comparable numbers of differentially abundant proteins were observed for both B6 and B6xD2 backgrounds when comparing 5XFAD to nontransgenic (Ntg) groups, demonstrating a similar overall effect of the transgene between mouse lines (**Supplemental Tables 5-6**). The overlap and concordance of AD genotype-driven protein changes between mouse lines were also compared (**Fig. 3**; **Supplemental Tables 7-8**). We identified a core group of proteins that were significantly changed in both mouse lines as well as protein subsets uniquely changed in only one mouse line or the other. Comparable findings were observed for both the cortex and hippocampal regions (**Supplemental Tables 7-10**). To understand the potential biological relevance of the changed proteins in each subset, gene ontology (GO) was performed in each brain region with attention to direction of change (**Fig. 3C-E**). Notably, “Amyloid-β binding” was among GO terms with the highest enrichment scores from proteins increased in both backgrounds in the cortex, indicating the consistency of core AD genotype-driven protein changes regardless of genetic background. The influence of genetic variation, however, also had clear impacts on disease relevant biological pathways. For example, the reactome term “Metabolism” was enriched in the opposite direction for each line, with B6 showing an increase and B6xD2 showing a decrease in proteins corresponding with this term. This is consistent with previously published work in which B6 and D2 mice exhibit significantly different glucoregulatory phenotypes (18). In addition, the directionality for some B6xD2 specific terms was unexpected in 5XFAD. For instance, the term “Aging” was decreased and the term “Neurotransmitter release” was increased on the B6xD2 line only. In summary, we found that the proportion of changed proteins with 5XFAD transgene expression on each line appeared comparable, and that key proteins affected by expression of the transgene were not different between B6 and B6xD2 lines. However, the impact of genetic background on AD genotype expression was significant for other proteins and pathways.

**Figure 3:**
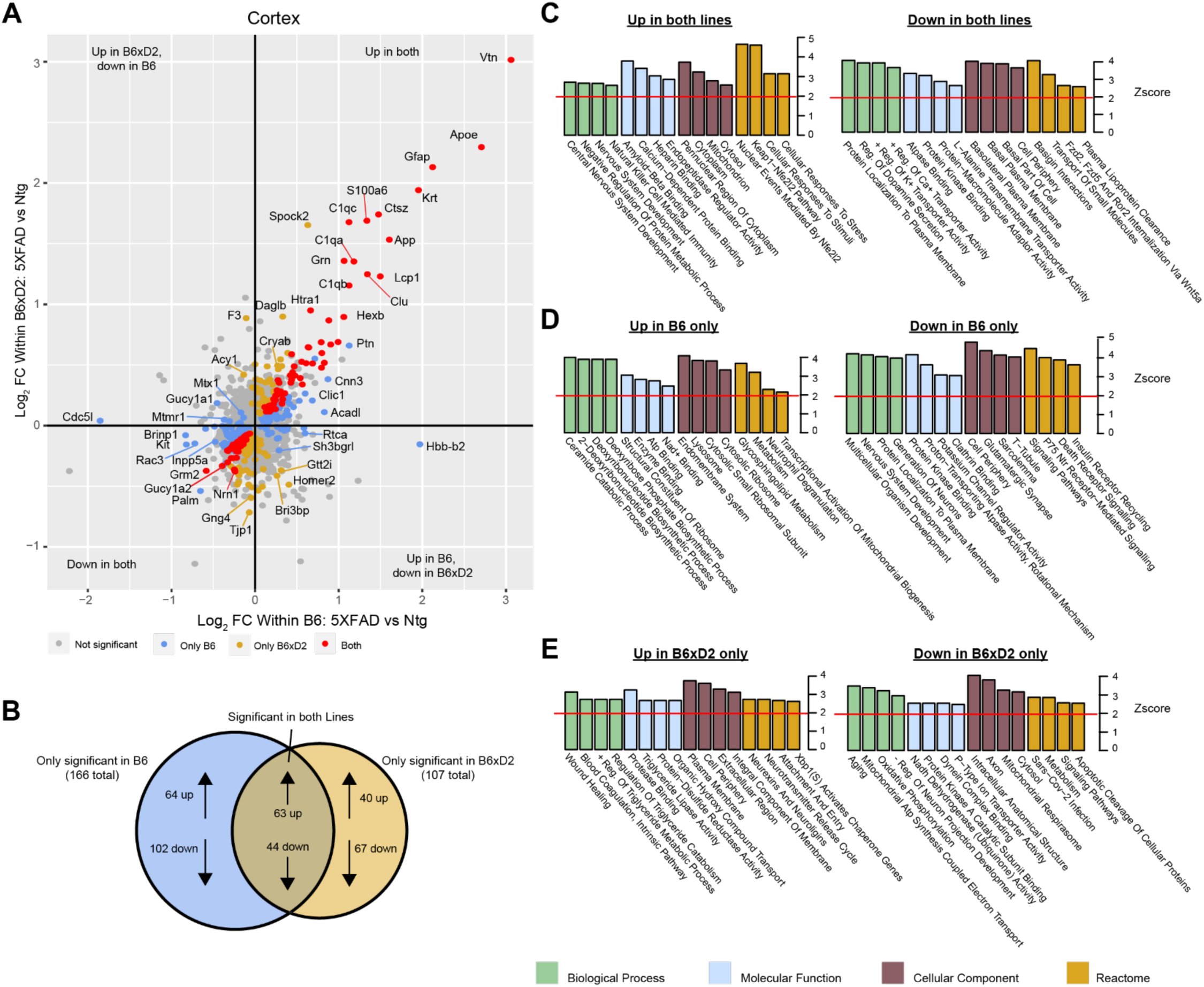
Differential protein abundance driven by genetic background and AD genotype. **A)** Comparison of differential protein abundance between 5XFAD and Ntg mice on either the B6 or B6xD2 genetic background in cortex. One-way ANOVA with Tukey post-hoc correction was used to generate p-values (significance was determined as p<0.05). Log^2^ fold change (FC) differences were plotted to incorporate directionality of change on each genetic background. Color scheme: gray, not significantly changed; red, significantly changed in both B6 and B6xD2; gold, significantly changed only in B6xD2; blue, significantly changed only in B6. **B)** Venn diagram representing overlap of significantly differential abundant proteins comparing 5XFAD with Ntg on either the B6 or B6xD2 genetic backgrounds (significance was determined as p<0.05). **C-E)** Gene ontology analysis of protein subsets significantly increased or decreased on both lines (**C**) or on only B6 (**D**) or only B6xD2 (**E**) lines. Ontology analysis was performed for biological process, molecular function, cellular component, and reactome. A Z-score of 1.96 or higher is considered significant (red line, *p*<0.05).

### Correlation network analysis of multi-region 5XFAD brain proteome

To examine biologically related groups of proteins and how they are altered by 5XFAD transgene expression and genetic background in hippocampus and cortex, we used the consensus Weighted Correlation Network Analysis (cWGCNA) algorithm (n=3,368 proteins) to build a protein co-expression network from our proteomic data. We identified 16 clusters (modules) of highly co-expressed proteins (**Fig. 4A; Supplemental Tables 11-12**). Modules were very highly preserved across both brain regions (**Fig. S2**). Representative module biology was assigned by top GO terms for module protein members (**Supplemental Table 13**). Only one module (M9) was assigned as ambiguous based on the available GO terms. Approximate cell type contribution to each module was determined by enrichment analysis of neuronal, microglial, astrocyte, oligodendrocyte and endothelial cell type markers (**Supplemental Tables 14-15**). There was at least one module significantly associated with each of the cell types.

**Figure 4:**
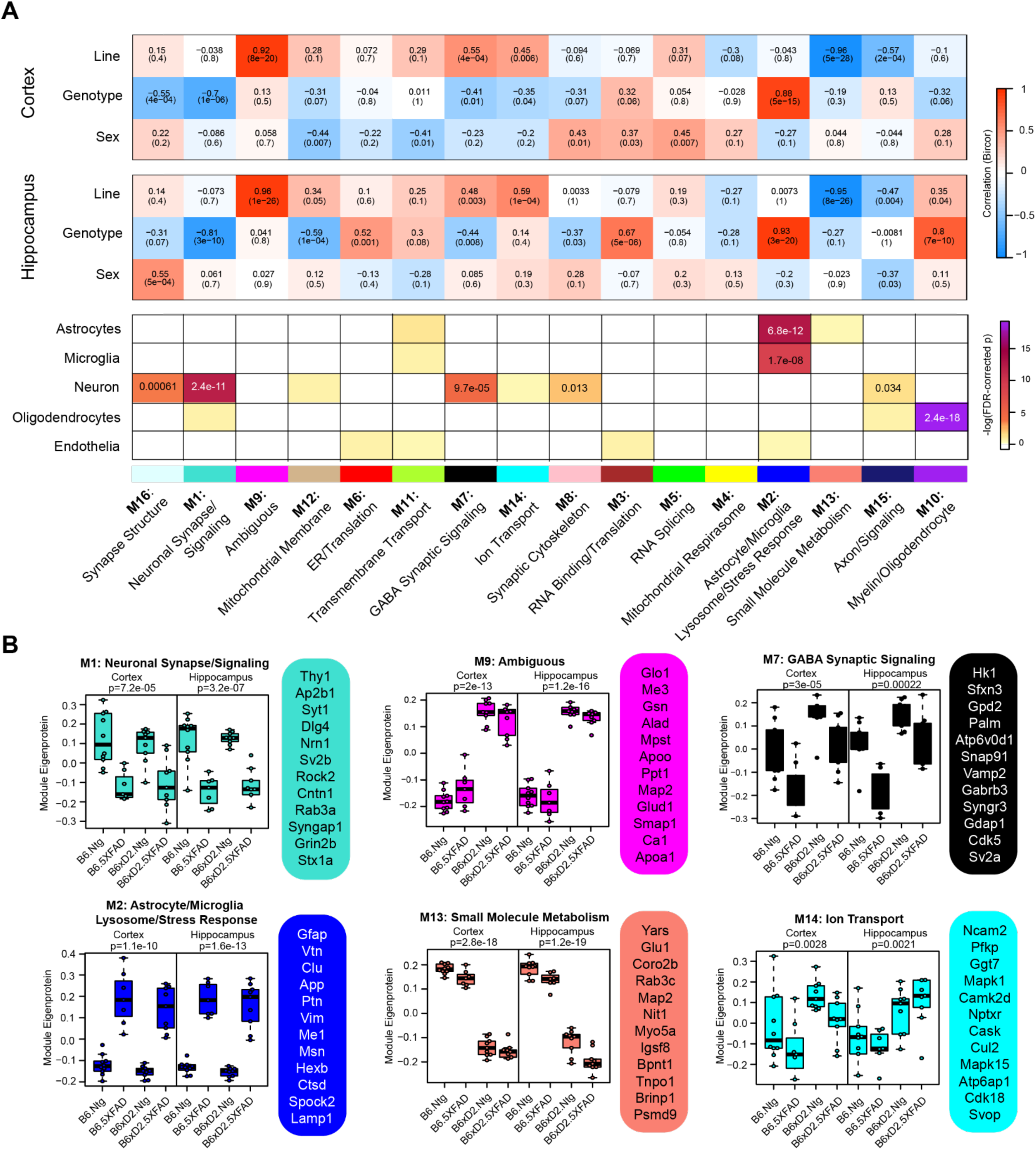
Consensus correlation network analysis of multi-region 5XFAD mouse brain proteome. **A)** <u>C</u>onsensus <u>w</u>ei<u>g</u>hted <u>c</u>orrelation <u>n</u>etwork <u>a</u>nalysis (cWGCNA) was performed with 3,368 proteins from cortex and hippocampus which resulted in 16 co-expression protein modules. (top) Correlation of module eigenproteins (MEs) with traits (line, AD genotype and sex) for each brain region is represented by heatmap (red, positive correlation; blue, negative correlation). (bottom) Enrichment of cell type markers in each module for astrocytes, microglia, neuron, oligodendrocytes and endothelial cell types. Modules are identified by color and number as well as representative biology as determined by top gene ontology terms. **B)** MEs for selected modules of interest grouped according to line—AD genotype pairings evaluated (B6 Ntg, B6 5XFAD, B6xD2 Ntg and B6xD2 5XFAD) in each brain region. Hub proteins of each module are provided beside each ME box plot. MEs were compared by group in each brain region using one-way ANOVA; unadjusted p-values are shown. Box plots represent median, 25^th^ and 75^th^ percentiles. Box hinges represent the interquartile range of the two middle quartiles with a group. Error bars are based on data points 1.5 times the interquartile range from the box hinge.

Module eigenproteins (MEs) were correlated with AD genotype, line and sex for each brain region. MEs represent the first principal component of protein expression of proteins within each module, and thus module— trait correlations can provide information on how variables of interest are related to these protein groups (**Fig. 4A**). We also examined ME levels across the different groups in hippocampus and cortex (**Fig. 4B**). Within the network, there were modules strongly driven by primary traits line and AD genotype as well as mixed effects. Modules M1 Neuronal Synapse/Signaling and M2 Astrocyte/Microglia Lysosome/Stress Response were highly related to AD genotype, where 5XFAD transgene expression was significantly associated with decreased protein abundance in M1 members and significantly associated with increased protein abundance in M2 module members, regardless of genetic background. Notably, M1 and M2 were the largest modules in the network (M1= 311 proteins, M2= 246 proteins) and represented the strongest cell-type enrichments for Neurons (M1) and astrocytes and microglia (M2). M9 Ambiguous and M13 Small Molecule Metabolism were the most significantly driven by genetic background, where M9 constituents were significantly higher in B6xD2 mice compared to B6 and M13 constituents were significantly lower in B6xD2 mice compared to B6, regardless of AD genotype. In addition, modules including M7 GABA Synaptic Signaling and M14 Ion Transport were significantly associated with more than one feature. Module M7 was significantly correlated with line and anticorrelated with AD genotype. Module M14 was correlated with line in both brain regions and anticorrelated with AD genotype only in the cortex.

In summary, large modules such as M1 Neuronal synapse/signaling and M2 Astrocyte/Microglia Lysosome/Stress Response indicate strong AD genotype driven effects related to disease unimpacted by genetic background. However, multiple modules strongly driven by line were also present. Together, these findings suggest that genetic background may have more modulatory effects on the proteomic changes surrounding amyloidosis observed in the 5XFAD model. Additional work is necessary to understand how the modules relating to genetic background may contribute to or impact the overall AD phenotype in these mice.

### Comparison of 5XFAD and human AD brain proteomic networks

In order to evaluate the extent to which genetic background influences model overlap with the human disease, we compared our mouse network with a human frontal cortex brain proteome network (16). We used two similar but distinct analyses to compare the mouse network with the human network: 1) module preservation, which assesses the presence or absence of mouse modules in the human network based on measures of network connectivity, and 2) overrepresentation analysis (ORA), which assesses enrichment of overlapping proteins across network modules. Module preservation analysis found that the majority of modules (10/16) in the mouse network were preserved with the human network, and 2 of these (M10 Oligodendrocyte/Myelin and M1 Neuronal Synapse/Signaling) were very strongly preserved (Fig. 5A). Both M10 and M1 were strongly correlated with AD genotype in both regions and to a lesser extent with line in the hippocampus (Fig. 4A). Modules M6 ER/Translation, M9 Ambiguous, M11 Transmembrane Transport, M12 Mitochondrial Membrane, M13 Small Molecule Metabolism and M14 Ion Transport were not preserved with the human network modules based on network connectivity. Of the mouse modules that did not preserve with human modules, M9, M13 and M14 in particular were strongly associated with genetic background. Overrepresentation analysis found greater overlap with human modules and identified modules in the mouse network that closely related to key human AD biology (Fig. 5B, **Supplemental Table 16**). All but 3 of the mouse network modules significantly overlapped with the human modules. Modules M11 Transmembrane Transport, M14 Ion transport and M13 Small Molecule Metabolism did not show significant overlap with any of the human modules, consistent with the module preservation analysis. Because our mouse cWGCNA network was a consensus network of hippocampal and cortex tissues, we also tested whether a cortex-only mouse network would show better preservation and overlap with the human cortex-only network (**Supplemental Table 17**). We found that 11/20 cortex-only modules were preserved in the human network, and 11/20 modules showed significant overlap by ORA, similar to the cWGCNA network findings (**Fig. S3**; **Supplemental Table 18**), indicating that the cross-species analyses were not significantly influenced by the inclusion of hippocampal tissue in the cWGCNA analyses. The collective results of these analyses indicate that the majority of protein co-expression relationships are preserved between mouse and human, but also suggest there may be significant effects driven by line that are either species specific or not well represented in this human network due to differences in proteome coverage.

**Figure 5:**
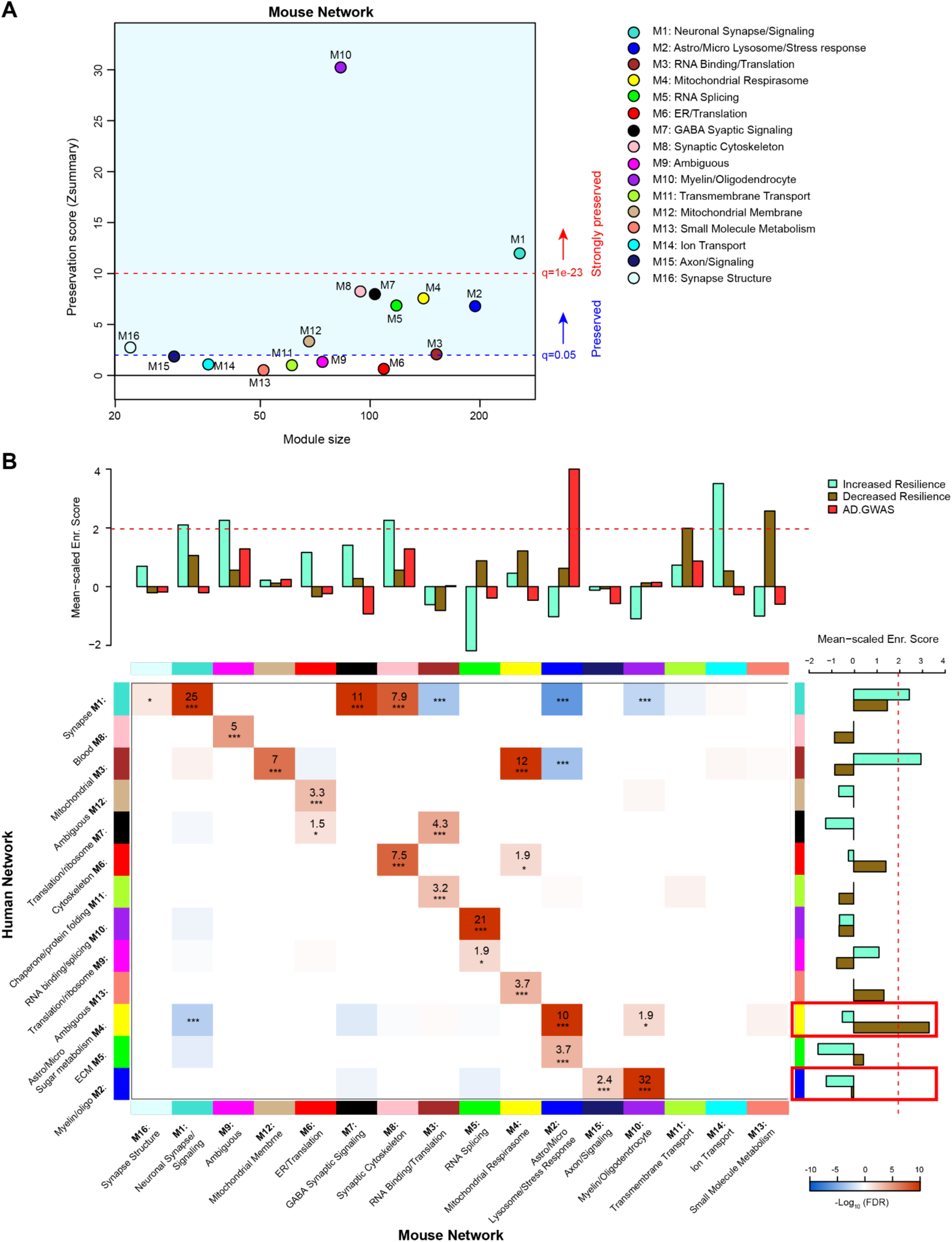
Comparison of 5XFAD mouse and human AD brain proteomic networks. **A)** Network preservation of mouse protein network modules in a human AD frontal cortex network as published in Johnson *et al.* (16). Modules with Zsummary greater than or equal to 1.96 (q=0.05, dashed blue line) are considered preserved, and modules with Zsummary of 10 or higher (q=1e-23, dashed red line) are considered highly preserved. The majority (10 out of 16) of consensus modules from the mouse proteome were found to be preserved in the human network. **B)** Heatmap for the overrepresentation analysis (ORA) of mouse consensus module members with human frontal cortex module members. Red indicates overrepresentation, blue indicates underrepresentation. Numbers in boxes are -log_10_ FDR values. *P* *<0.05, **<0.01, ***<0.001. Heatmap threshold is set at 10% FDR (0.1). Bar plots in heatmap margins show enrichment of proteins identified from GWAS of AD risk (red) or PWAS of cognitive resilience for both increased resilience (teal) or decreased resilience (olive) for mouse and human network module members. Dashed red line (z-score 1.96 or FDR q=<0.05) indicates significance cutoff. Red boxes in human enrichment bar plot indicate previously identified significantly enriched modules for AD GWAS proteins.

To assess how the mouse network modules related to human AD genetic risk, an enrichment analysis was performed using results from AD genome wide association studies (GWASs) (19–21). This enrichment strategy previously identified human modules M2 Myelin/oligodendrocytes and M4 Astrocyte/Microglia sugar metabolism as enriched with AD risk factor proteins (16). The same analysis was performed on mouse modules, which identified M2 Astrocyte/Microglia Lysosome/Stress Response as enriched with AD risk factor proteins (**Supplemental Table 19**). Importantly, ORA results indicated mouse module M2 Astrocyte/Microglia Lysosome/Stress Response significantly overlapped with the human module M4 Astrocyte/Microglia Sugar metabolism. This finding highlights the consistency of human disease features represented in the mouse model. Also of note, mouse module M2 was driven strongly by AD genotype but not genetic background.

In addition to disease risk, we also used results from a proteome wide association study (PWAS) of cognitive resilience to identify modules in both networks enriched with proteins shown to relate to cognitive performance over time (22). PWAS results were generated from dorsolateral prefrontal cortex brain tissue samples analyzed via tandem mass tag (TMT) -MS which were adjusted for AD pathologies before determining the association between cortical protein abundance with cognitive change over time. Proteins with increased abundance and a relationship to slower rates of cognitive decline were considered to confer increased resilience whereas proteins with higher abundance and associated with faster rates of decline were considered to confer decreased resilience. Multiple modules were identified in both human and mouse networks for increased and decreased resilience proteins. There were four mouse modules enriched for proteins associated with increased cognitive resilience: M1 Neuronal synapse/signaling, M9 Ambiguous, M8 Synaptic cytoskeleton and M14 Ion transport (**Supplemental Table 20**). There were also two mouse modules enriched for proteins associated with decreased resilience: M11 Transmembrane transport and M13 Small molecule metabolism (**Supplemental Table 21**). In the human network, there were two modules enriched for increased resilience proteins: M1 Synapse and M3 Mitochondrial (**Supplemental Table 22**). Only one module was enriched in the human network for decreased resilience proteins: M4 Astrocyte/Microglia Sugar metabolism (**Supplemental Table 23**).

In summary, consistency in enrichment results could be observed between networks, however, there were also distinct, non-overlapping features. Mouse module M2 Astrocyte/Microglia Lysosome/Stress Response, which was enriched for AD GWAS markers, most strongly overlapped with human module M4 Astrocyte/Microglia Sugar metabolism, which was the top GWAS module in the human network (16). Mouse modules M1 Neuronal synapse/signaling, M9 ambiguous and M8 Synaptic cytoskeleton were all enriched with increased resilience markers and strongly overlapped with human module M1 Synapse, which was consistently enriched with increased resilience markers. Notably, mouse modules M13 Small molecule metabolism, M11 Transmembrane transport and M14 Ion transport all significantly enriched for either increased or decreased resilience protein markers, however, these modules had no significant overlap with human modules.

## Discussion

In this study, we examined the brain proteomes of mice from two genetic backgrounds across two brain regions of 5XFAD and Ntg animals to assess the impact of incorporating genetic diversity on the proteome in mouse models of AD. We found that both genetic background and transgene expression contributed strongly to variance in the proteome. The strongest protein changes driven by the 5XFAD transgene as reflected in the M2 Astrocyte/Microglia Lysosome/Stress Response and M1 Neuronal Synapse/Signaling modules were not significantly altered by genetic background at either the single protein or protein network levels. However, genetic background did influence proteins relevant to disease related to GABA synaptic signaling biology and mitochondrial processes, among others. Together, these results indicate that, as expected, the classical disease-relevant changes related to 5XFAD transgene expression are robust; however, proteins significantly correlated to genetic background may represent individualized targets related to novel genetic factors and disease subtypes, and may also provide insights into resilient and susceptible phenotypes. Network analysis also facilitated comparison of mouse and human co-expression brain proteomes, in which the majority of the mouse modules preserved and overlapped with human network modules. Modules that were strongly affected by AD in the human network overlapped with modules that were most strongly driven by the 5XFAD transgene in the mouse network. Overall, this study contributes to ongoing efforts to rigorously validate and characterize AD model systems using multiple different –omic approaches, and illustrates the potential effects on the mouse proteome of introducing genetic variability for improving cross-species translation.

Previous studies have documented line-dependent changes in pathological and phenotypic measures, including Aβ deposition and learning and memory tasks (6, 10). Consistently, the brain proteome comparing 5XFAD with Ntg animals demonstrated line-specific differences at the individual protein and co-expression levels. Among these line-specific differences, biology relating to synaptic function could suggest potentially important changes introduced by genetic variability relevant to human neurodegenerative conditions. Specifically, ontology terms “Neurotransmitter release cycle” and “Neurexins and neuroligins” were significantly changed in B6xD2 animals. In addition, module M7 GABA Synaptic Signaling was strongly correlated with line. Changes in synaptic integrity and density are believed to underly cognitive symptoms in human disease and more recently the ability to maintain functional synaptic connections has been proposed as a mechanism of supporting cognitive resilience in AD (23, 24). Inherent differences in synaptic biology introduced by genetic variability in mouse models of AD could represent a novel opportunity to investigate human relevant alterations in neuronal communication. The extent to which genetic diversity affects synaptic changes requires further study.

One important aspect of this study is the inclusion of more than one brain region for analysis. We found that the hippocampus was more strongly affected by transgene expression than cortex. Principal component and variance partition analyses demonstrated both AD genotype and line strongly contribute to variance in the proteome. However, the strength of this association was muted when the two brain regions were not separated, in which region became the strongest explanation of variance in the model (data not shown). Consensus network analysis also identified multiple modules strongly influenced by brain region including module M10 Myelin/oligodendrocyte, in which correlation with AD genotype was inverse by region. Collectively these findings indicate the strength of evaluating multiple brain regions when validating specific models.

Previous methods of introducing genetic complexity in AD mouse models include the use of wild-derived lines (25). Genetically diverse, wild-derived transgenic mice produced strong differentiation of phenotypes associated with important human disease features including cognitive-like changes, neurodegeneration and amyloid dynamics. However, this method of introducing genetic diversity has been cautioned due to the lack of rigorous characterization in the absence of AD transgenes (3). The use of fully inbred parental lines, such as B6 and D2 that have been extensively characterized, overcomes this ambiguity. Therefore, the genetic variability introduced by crossing the B6 and D2 lines would allow differentiation of highly heritable phenotypes and traits relating to risk and resilience in AD progression (13). Moreover, this strategy is not limited specifically to the B6 and D2 lines, but rather an adaptable reference in the utility of this strategy for future model development and characterization. Taken together, introducing genetic complexity by crossing widely used inbred mice represents a potentially advantageous research tool for both improving recapitulation of complex human phenotypes and practical reproducibility.

Comparing the mouse and human networks highlighted important findings: 1) mouse modules most strongly representative of human disease were driven by 5XFAD transgene overexpression and not significantly influenced by genetic background, and 2) certain mouse modules did not overlap with the human network at all. Mouse module M2 Astrocyte/Microglia Lysosome/Stress Response enriched for AD GWAS markers and corresponded most significantly to the human module M4 Astrocyte/Microglia Sugar Metabolism, which also enriched for AD GWAS markers. This overlap of disease-relevant modules enriched for glial cells is well supported in the literature (26). Similarly, mouse module M1 Neuronal Synapse/Signaling overlapped significantly with human module M1 Synapse, both of which enriched for markers of increased cognitive resilience. These mouse modules (M2 Astrocyte/Microglia Lysosome/Stress Response and M1 Neuronal Synapse/Signaling) were significantly correlated to AD genotype, but not genetic background. These findings suggest that in this model there are core pathologies and features at the proteomic level related to human disease that are strongly driven by 5XFAD transgene expression, as expected. They also suggest other modules that are strongly driven by genetic background could provide putative targets for understanding individual differences driven by genetic variants that may inform mechanisms of resilience or susceptibility. In addition, the lack of overlap of specific mouse modules (M11 Transmembrane Transport, M14 Ion Transport and M13 Small Molecule Metabolism) might also be explained by the current transgenic model utilized. Another consideration for this gap is the comparative depth of LFQ versus TMT proteomics. Both mouse and human networks compared here were analyzed via LFQ proteomics, however, the PWAS results integrated in the enrichment analysis were generated using TMT-MS. This offers a potential explanation for the significant enrichment of M11, M14 and M13 with proteins associated with cognitive trajectory despite their lack of overlap with human modules. Future proteomic studies on APP knock-in models using TMT-MS to increase proteome depth of coverage may allow for more sensitive detection of genetic background effects on disease-relevant proteomic changes.

Some potential limitations of our study should be noted. First, the current cohort of animals were aged 12-14 months, a timepoint in which significant amyloid pathology has developed in the 5XFAD model. While evaluating genetic contribution is valuable at this stage in the lifespan, measuring proteomic differences at one static timepoint may miss transient or time-dependent pathological changes. Future studies evaluating multiple timepoints could, therefore, provide valuable insights into rates of proteomic changes that may be influenced by mixed genetic backgrounds. Second, one of the goals of introducing genetic complexity in mouse models is to generate phenotypically distinct substrains that enable characterization of novel genetic variants contributing to variable trajectories of cognitive impairment. Here we analyzed only one mixed genetic background (B6xD2), and behavioral data were not included in the analysis. Analysis of larger cohorts of up to 50 AD-BXD lines that better model heterogeneity of humans are underway that will provide sufficient power to interpret the mediation of the proteome on diverse brain and cognitive phenotypes (13, 27). These rich data are being generated and shared from consortium initiatives Resilience-AD and MODEL-AD (Model Organism Development and Evaluation for Late-Onset Alzheimer’s Disease) (7).

Mice represent an ideal model system due to their combination of phylogenetic conservation with humans and their relative ease in experimental manipulation and management. Currently, there are nearly 200 (and counting) transgenic models of AD that have been developed, the majority of which utilize FAD mutations expressed on the fully inbred B6 genetic background. This schema for model development has led to invaluable insights into the core mechanisms of AD progression and risk, but clearly there remains a need for improvement in translational validity of AD mouse models. Proteomics provides one level of analysis into a complex, polygenic human disease. Incorporation and careful consideration of other levels of analysis, such as the transcriptome (26), can provide additional insights into the effect of genetic variability on AD phenotypes for advancement of both model translatability and therapeutic target nomination. Our results suggest that 5XFAD transgene expression induces robust changes on the mouse brain proteome but that the addition of genetic background complexity introduces significant proteomic variance that may contribute to phenotypic variability and alignment to the human disease. Therefore, this approach remains a promising avenue to improve face validity in other AD mouse models such as knock-in models where proteomic changes may be more subtle than transgenic models.

## Supporting information

Hurst et al Supplementary Tables

## Data and Code availability

Raw mass spectrometry data and database search results from cortex and hippocampus mouse brain tissue analysis can be found at https://www.synapse.org/B6xD2proteomics. Processed data and code are also provided. Human MS data was previously shared and can be accessed at https://www.synapse.org/consensus.

## Author Contribution Statement

CH, AD, CK, and EJ designed the experiments; CH, AD, and DD carried out experiments; CH, EBD, DD, and EJ analyzed data; AD, NTS, and CK provided advice on the interpretation of data and manuscript review; CH and EJ wrote the manuscript with input from coauthors.

## Acknowledgements

This study was supported by AARF18565506 (AD), R01AG061800 (NS), R01AG057914 (CK), R01AG054180 (CK), R01AG075818 (CK), RF1AG059778 (CK), and K08AG068604 (EJ).

## Methods and Materials

### Mice

Female hemizygous 5XFAD mice on a congenic C57BL/6J background (RRID: MMRC_034848-JAX) were bred to male C57BL/6J (RRID: IMSR_JAX:000664) or DBA/2J mice (with corrected *Gpnmb* mutation, RRID: IMSR_JAX:007048). Animals were kept on a 12:12 light:dark cycle and were provided food and water ad libitum. All routine procedures were approved by the Institutional Care and Use Committee (IACUC) at The Jackson Laboratory, and in accordance with the standards of the Association for the Assessment and Accreditation of Laboratory Animal Care (AAALAC) and the National Institutes of Health Guide of the Care and Use of Laboratory Animals. Male and female mice (balanced across groups) at 14 mo of age (+/- 2 wks), animals were deeply anaesthetized with isoflurane and rapidly decapitated and the brain removed. Hippocampus and frontal cortices were isolated on an ice-cold dissecting block and immediately frozen in liquid nitrogen.

### Tissue homogenization

Tissue dissections of cortex and hippocampus were each homogenized in urea lysis buffer (8 M urea, 100 mM NaH2PO4, pH8.5) at a 1:5 weight to buffer ratio with HALT phosphatase and protease inhibitor cocktail (1X final concentration, Pierce). Samples were homogenized in RINO sample tubes (Next Advance) with ∼100 ul of stainless-steel beads using a Bullet Blender (Next Advance) at 4° for two full 5-minute cycles. Homogenates were transferred to clean, Eppendorf LoBind tubes and sonicated for three cycles at 5 seconds on and 5 seconds off at 20% amplitude, on ice. Samples were then centrifuged for 5 minutes at 15,000xg and supernatant transferred to clean tubes. Protein concentrations were determined by bicinchoninic acid assay (BCA, Pierce). One dimensional SDS-PAGE gels were run followed by Coomassie blue staining as quality control to ensure protein integrity and equal loading.

### SDS-PAGE

20 ug of protein from each sample was mixed with Laemmli sample buffer (BioRad) and β-mercaptoethanol before being boiled at 95° for 10 minutes, spun briefly to collect volume and loaded into Bolt 4-12% Bis-Tris gradient gels (Invitrogen). Loaded gels were initially electrophoresed at 80 mV for the lowest percentage gradient of the gel followed by 120 mV for the remainder of the gel. Gels were then submerged in Coomassie blue staining overnight and destained briefly the following day to visualize protein banding (**Supplemental Figure 4**).

### Protein digestion

100 ug of protein from each sample was aliquoted and volume normalized to 50 uL in urea lysis buffer before being reduced with 5 mM dithiothreitol (DTT) for 30 minutes at room temperature followed by alkylation with 10 mM iodoacetamide (IAA) for 30 minutes, light protected. Samples were digested overnight in 1:50 (w/w) lysyl endopeptidase (Wako). The following day, urea concentration for each sample was diluted to <1 M using ammonium bicarbonate (ABC) and digested for an additional ∼16 hours using 1:50 (w/w) trypsin (Promega). Following digestion, peptides were acidified to a final concentration of 1% (v/v) formic acid (FA) and 0.1% trifluoronic acid (TFA) and desalted using 30 mg HLB columns (Oasis). Prior to sample loading, each HLB column was rinsed with 1 mL of methanol, washed with 1 mL 50% (v/v) acetonitrile (ACN) and equilibrated (x2) with 1 mL 0.1% (v/v) TFA. Samples were then loaded onto the columns and washed (x2) with 1 mL 0.1% (v/v) TFA. Peptides were eluted with 2 volumes of 0.5 mL 50% (v/v) ACN. Eluates were frozen in -80° C overnight before being completely dried using a SpeedVac (LabConco).

### LC-MS/MS

All samples (∼1ug) were loaded and eluted by an Ultimate RSLCnano (Thermofisher Scientific) with an in-house packed 15 cm, 150 μm i.d. capillary column with 1.7 μm C18 CSH (Waters) over a 45 min gradient. The gradient went from 1 to 99% Buffer B (Buffer A: water in 0.1% formic acid and Buffer B: 80% acetonitrile in 0.1% formic acid). Mass spectrometry was performed with an Orbitrap Lumos (Thermo) in positive ion mode using data-dependent acquisition with 3 second top speed cycles. Each cycle consisted of one full MS scan followed by as many MS/MS events that could fit within the given 1 second cycle time limit. MS scans were collected at a resolution of 120,000 (375-1500 m/z range, 4x10^5 AGC, 50 ms maximum ion injection time). Only precursors with charge states between 2+ and 6+ were selected for MS/MS. All higher energy collision-induced dissociation (HCD) MS/MS spectra were acquired at a resolution of 15,000 (1.6 m/z isolation width, 35% collision energy, 1×10^5 AGC target, 22 ms maximum ion time). Dynamic exclusion was set to exclude previously sequenced peaks for 30 seconds within a 10-ppm isolation window.

### Database searching and protein quantification

Quantitation was performed as previously published (28), with slight modification: RAW data files from all 80 samples were analyzed using MaxQuant (v1.6.17.0) using a mouse database (91,415 target sequences, downloaded August 2020). Methionine oxidation, asparagine and glutamine deamidation, protein N-terminal acetylation, and serine, threonine and tyrosine phosphorylation were variable modifications (up to 5 allowed per peptide); cysteine was assigned as a fixed carbamidomethyl modification (+57.0215 Da). A precursor mass tolerance of ±20 ppm was applied prior to mass accuracy calibration and ±4.5 ppm after internal MaxQuant calibration. Other search settings included a maximum peptide mass of 6,000 Da, a minimum peptide length of 6 residues, and 0.05 Da tolerance for high resolution MS/MS scans. The false discovery rate (FDR) for peptide spectral matches, proteins, and site decoy fraction were all set to 1 percent. Quantification settings were as follows: re-quantify with a second peak finding attempt after protein identification has completed; match full MS1 peaks between runs; a 0.7 min retention time match window was used after an alignment function was found with a 20-minute retention time search space. Only razor and unique peptides were considered for protein level quantitation as summed intensities.

### Data preprocessing

A total of 4,351 high confidence, master proteins were identified and quantified across all 80 samples. Protein abundances were log2 transformed before sample outlier detection and missingness filtering. Network connectivity identified 5 samples as outliers (samples greater than 3 standard deviations away from the mean, as previously described (14, 16)) that were removed along with the matched tissue pairs from the other brain region and only proteins with >50% missing values across samples were included. The final data matrix included 70 samples and 3,368 proteins for downstream analyses.

### Differential expression analysis

Significantly differentially changed proteins between groups were defined using one-way ANOVA with Tukey’s comparison post hoc test (significance was determined as p<0.05). Differential expression displayed as volcano plots were generated using the ggplot2 package (3.3.5).

### Consensus Weighted Gene Correlation Network Analysis (cWGCNA)

Network analysis was performed using the consensus configuration of the Weighted Gene Correlation Network Analysis (cWGCNA, version 1.69) algorithm to identify co-expression modules present and shared in both cortex and hippocampal brain regions. The WGCNA::blockwiseConsensusModules function was run with soft threshold power at 7.0, deepsplit of 4, minimum module size of 30, merge cut height at 0.07, mean topological overlap matrix (TOM) denominator, using bicor correlation, signed network type, pamStage and pamRespectsDendro parameters both set to TRUE and a reassignment threshold of 0.05. This function calculates pairwise biweight mid-correlations (bicor) between protein pairs. The resulting correlation matrix is then transformed into a signed adjacency matrix which is used to calculate a topological overlap matrix (TOM), representing expression similarity across samples for all proteins in the network. This approach uses hierarchical clustering analysis as 1 minus TOM and dynamic tree cutting lends to module identification. Following construction, module eigenprotein (ME) values were defined. The MEs are the first principle component of a given module and are considered representative abundance values for a module that also explain modular protein covariance (29). Pearson correlation between proteins and MEs was used as a module membership measure, defined as kME.

### Gene Ontology (GO) and cell type marker enrichment analyses

Gene ontology (GO), Wikipathway, Reactome and molecular signatures database (MSigDB) term enrichment in our gene sets of mouse network module members and significantly differentially changed protein subsets was determined by gene set enrichment analysis (GSEA) using an in-house developed R script (https://github.com/edammer/GOparallel). Briefly, this script performs one tailed Fisher’s exact tests (FET) enabled by functions of the R piano package for ontology enrichment analysis on gene sets downloaded from http://baderlab.org/GeneSets, which is maintained and updated monthly to pull in current gene sets from more than 10 different database sources including those mentioned above (30, 31). Redundant core GO terms were pruned in the GOparallel function using the minimal_set function of the ontologyIndex R package (32). Cell type enrichment was also investigated as previously published (14, 28, 33). An in-house marker list combined previously published cell type marker lists from Sharma *et al.* (34) and Zhang *et al.* (35) were used for the cell type marker enrichment analysis for each of the five cell types assessed (neuron, astrocyte, microglia, oligodendrocyte and endothelial). If, after the lists from Sharma *et al.* and Zhang *et al.* were merged, gene symbol was assigned to two cell types, we defaulted to the cell type defined by the Zhang et al. list such that each gene symbol was affiliated with only one cell type. Fisher’s exact tests were performed using the cell type marker lists to determine cell type enrichment and were corrected by the Benjamini-Hochberg procedure.

### Network Preservation

Network preservation was determined using the WGCNA::modulePreservation() function. Zsummary composite preservation scores were calculated using the mouse network as the test network and the human network and the reference network. Parameters included: 500 permutations, random seed set to 1 (for reproducibility) and quickCor was set to 0.

### Enrichment analysis of Proteome Wide Association Study (PWAS) and Genome Wide Association Study (GWAS) results

The proteome Wide Association Study (PWAS) of cognitive trajectory by Yu *et al*. tested 8,356 proteins for correlation to change in cognition over time (22). Unique gene symbols representing protein gene products positively correlated (n=645) and negatively correlated (n=575) to cognitive resilience were split into lists with corresponding *P* values. For GWAS of AD risk, compiled single nucleotide polymorphism (SNP) summary statistics were used ((19–21), <u>MAGMA.SPA/MAGMAinput.zip at main · edammer/MAGMA.SPA · GitHub</u>). These lists were separately checked for enrichment in both mouse and human network modules using a permutation-based test (10,000 permutations) implemented in R with exact *P* values for the permutation tests calculated using the permp function of the statmod package (1.4.36). Module-specific mean *P* values for enrichment were determined as a *Z* score, specifically as the difference in mean *P* value of gene product proteins hitting a module at the level of gene symbol minus the mean *P* value of genes hit in the 10,000 random replacement permutations, divided by the standard deviation of *P* value means also determined in the random permutations (**Supplemental Tables 19-23**).

### Additional statistical analyses

All proteomic statistical analyses were performed in R (version 4.0.3). Box plots represent the median and 25^th^ and 75^th^ percentile extremes; the hinges of a box represent the interquartile range of the two middle quartiles of data within a group. Error bars extents are defined by the farthest data points up to 1.5 times the interquartile range away from the box hinges. Correlations were performed using biweight midcorrelation function from the WGCNA package.

---

**Supplemental Figure 1:**
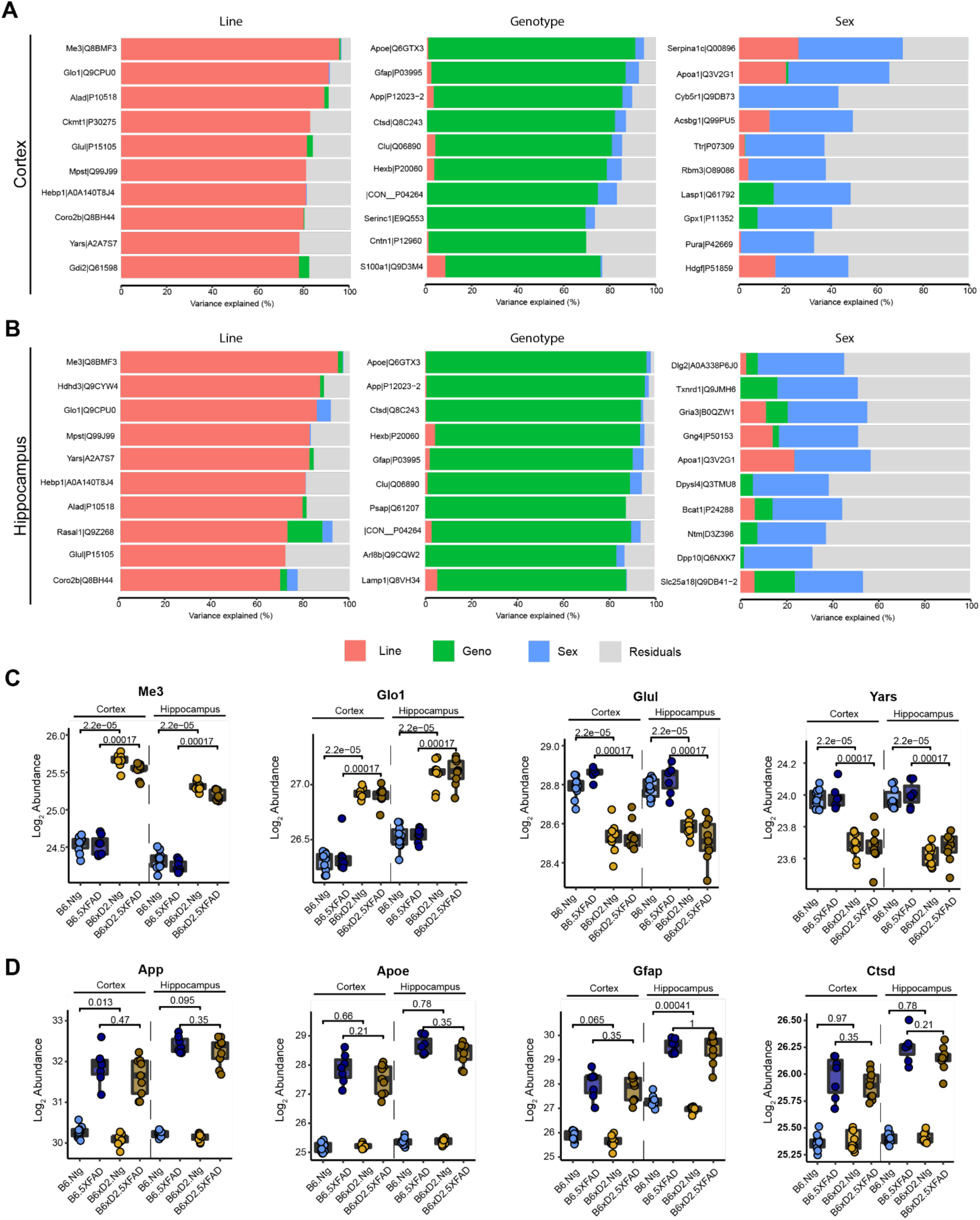
Stacked bar plots illustrating the top 10 proteins explaining greatest variance for line, AD genotype and sex in cortex (**A**) and hippocampus (**B**). **C)** Top variance explained proteins for line across cortex and hippocampus. **D)** Top variance explained proteins for AD genotype across cortex and hippocampus.

**Supplemental Figure 2:**
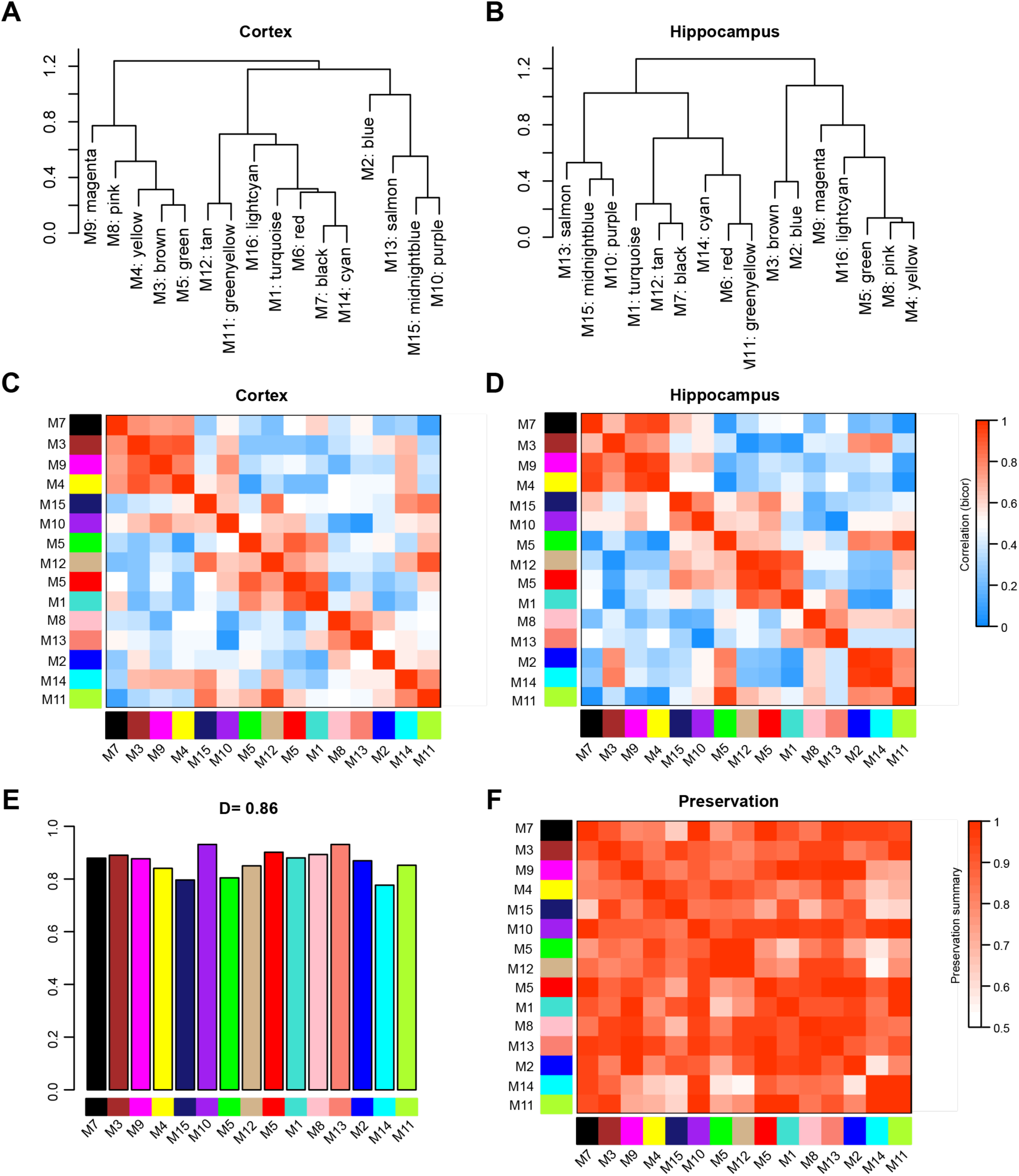
**A-B**) Consensus module eigneprotein clustering represented as a dendrogram for cortex (**A**) and hippocampus (**B**). **C-D**) Module eigenprotein correlation heatmap for cortex (**C**) and hippocampus (**D**). **E)** Mean preservation relationship for each eigenprotein was calculated for the consensus network. Mean preservation was 0.83, indicating a high degree of preservation. **F)** Preservation adjacency of the consensus network represented by heatmap. Most relationships were highly preserved across brain regions.

**Supplemental Figure 3:**
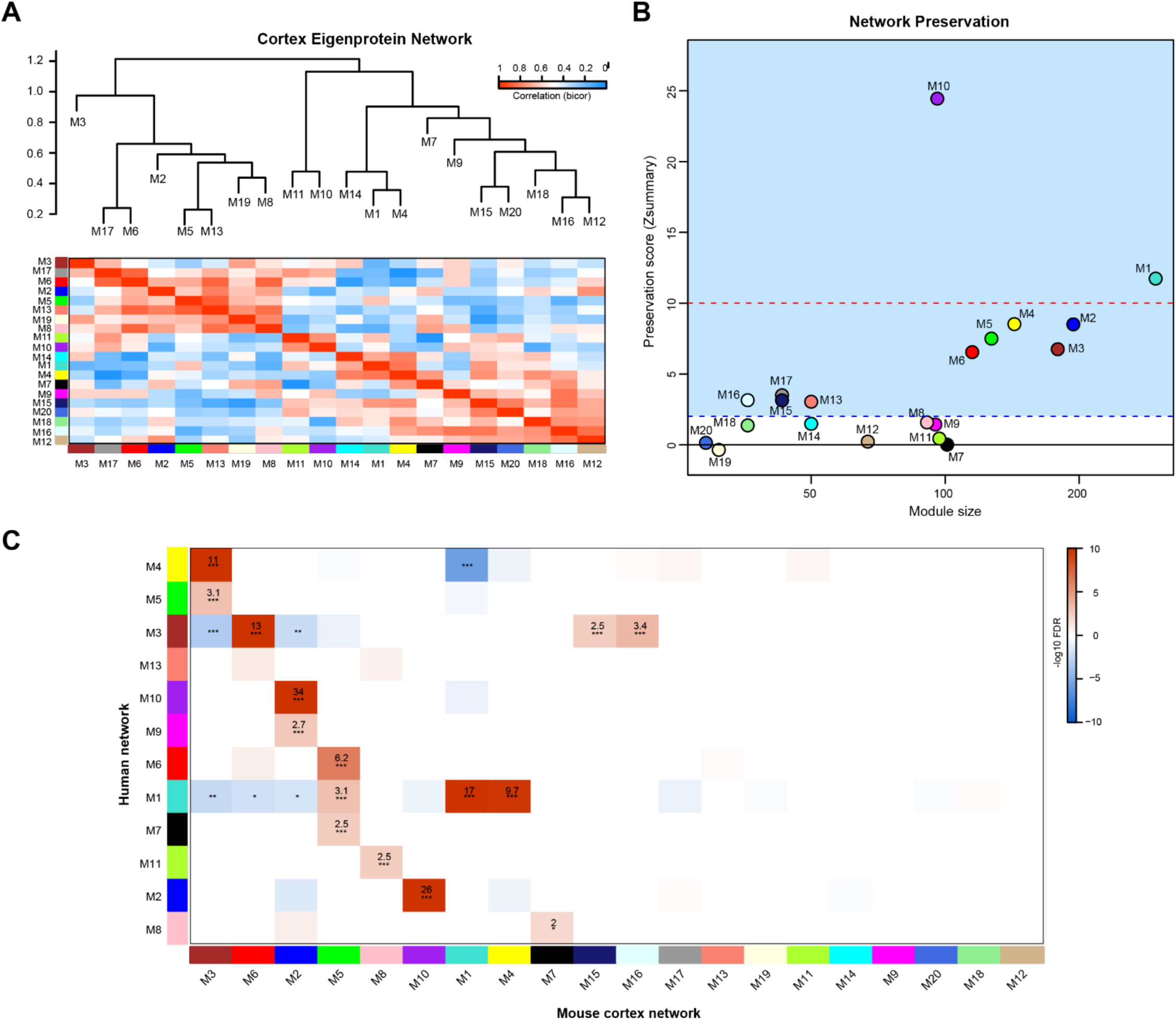
**A)** Eigenprotein network built using only cortex data showing module relatedness dendrogram and correlation (bicor). **B)** Network preservation of mouse (cortex only) protein network in human frontal cortex network (16). Modules with Zsummary greater than or equal to 1.96 (q=0.05, dashed blue line) are considered preserved, and modules with Zsummary of 10 or higher (q=1e-23, dashed red line) are considered highly preserved. The majority (11 out of 20) of consensus modules from the mouse (cortex only) proteome were found to be preserved in the human network. **C)** Heatmap for the overrepresentation analysis (ORA) of mouse (cortex only) module members with human frontal cortex module members. Numbers in boxes are -log_10_ FDR values. *P* *<0.05, **<0.01, ***<0.001. Heatmap threshold is set at 10% FDR (0.1).

**Supplemental Figure 4:**
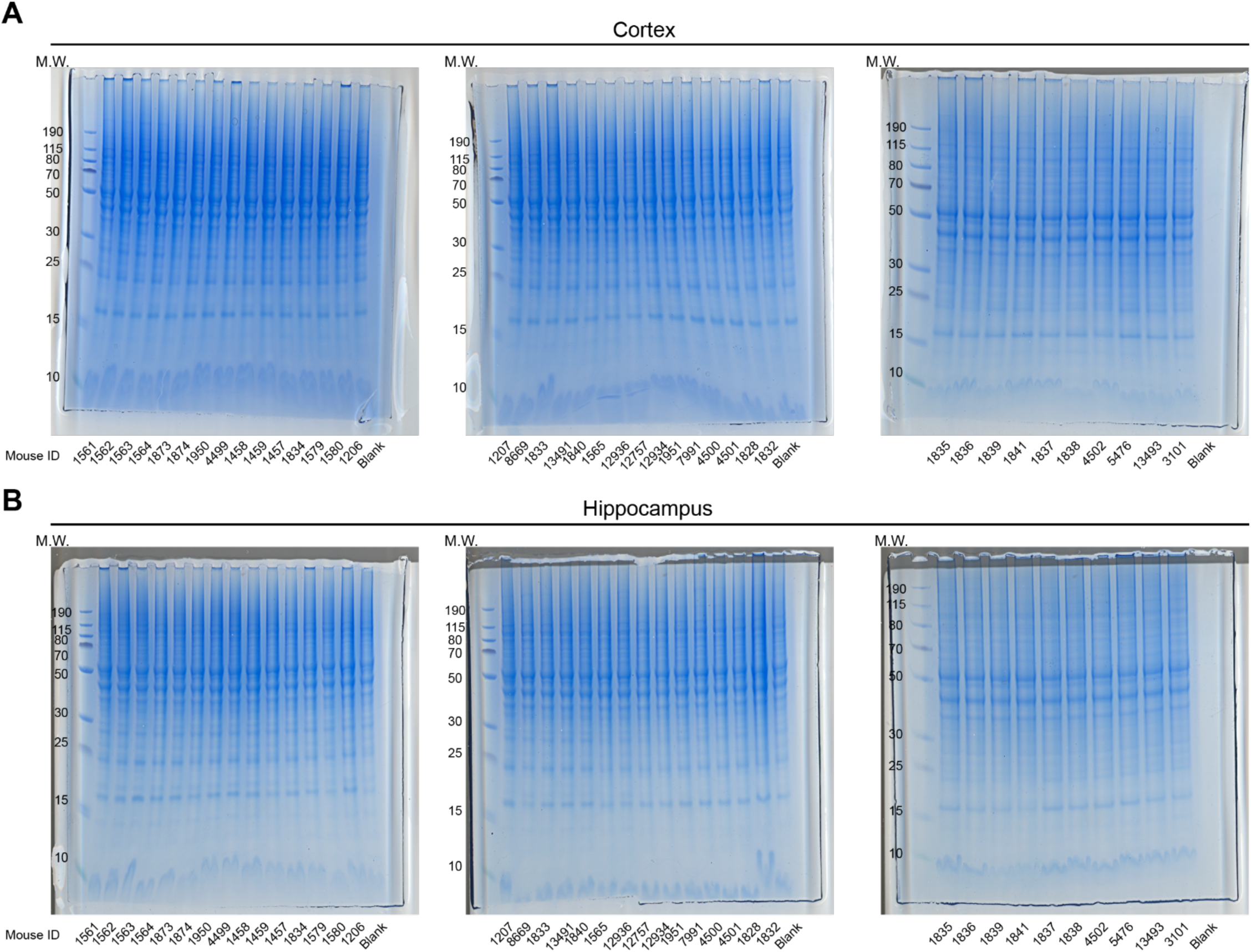
**A-B**) Protein integrity was assessed by SDS-PAGE gel stained with Coomassie blue in the Cortex (**A**) and Hippocampus (**B**).

